# Rootstocks shape the rhizobiome: Rhizosphere and endosphere bacterial communities in the grafted tomato system

**DOI:** 10.1101/375444

**Authors:** Ravin Poudel, Ari Jumpponen, Megan M. Kennelly, Cary L. Rivard, Lorena Gomez-Montano, Karen A. Garrett

## Abstract

Root-associated microbes are critical to plant health and performance, although understanding of the factors that structure these microbial communities and theory to predict microbial assemblages are still limited. Here we use a grafted tomato system to study the effects of rootstock genotypes and grafting in endosphere and rhizosphere microbiomes that were evaluated by sequencing 16S rRNA. We compared the microbiomes of nongrafted tomato cultivar BHN589, selfgrafted BHN589, and BHN589 grafted to Maxifort or RST-04-106 hybrid rootstocks. OTU-based bacterial diversity was greater in Maxifort compared to nongraft controls, whereas bacterial diversity in the controls (selfgraft and nongraft) and the other rootstock (RST-04-106) was similar. Grafting itself did not affect bacterial diversity; diversity in the selfgraft was similar to the nongraft. Bacterial diversity was higher in the rhizosphere than in the endosphere for all treatments. However, despite the lower overall diversity, there was a greater number of differentially abundant OTUs (DAOTUs) in the endosphere, with the greatest number of DAOTUs associated with Maxifort. In a PERMANOVA analysis, there was evidence for an effect of rootstock genotype on bacterial communities. The endosphere-rhizosphere compartment and study site explained a high percentage of the differences among bacterial communities. Further analyses identified OTUs responsive to rootstock genotypes in both the endosphere and the rhizosphere. Our findings highlight the effects of rootstocks on bacterial diversity and composition. The influence of rootstock and plant compartment on microbial communities indicates opportunities for the development of designer communities and microbiome-based breeding to improve future crop production.

**Importance:** Understanding factors that control microbial communities is essential for designing and supporting microbiome-based agriculture. In this study, we used a grafted tomato system to study the effect of rootstock genotypes and grafting on bacterial communities colonizing the endosphere and the rhizosphere. Comparing the bacterial communities in control treatments (nongraft and selfgraft plants) with the hybrid rootstocks used by farmers, we evaluated the effect of rootstocks on overall bacterial diversity and composition. These findings indicate the potential for using plant genotype to indirectly select bacterial taxa. In addition, we identify taxa responsive to each rootstock treatments, which may represent candidate taxa useful for biocontrol and in biofertilizers.

## Introduction

The root-associated microbiome, or “rhizobiome”, is essential for plant health and performance (1, 2). Some microbes in the rhizobiome are especially important for plants during harsh and unfavorable growing conditions (3). Microbe-mediated nutrient uptake, disease resistance, and stress tolerance (1, 4, 5) are some examples of microbial functions important to agriculture and host biology more generally. While the importance of microbes in provisioning ecosystem services is clear, greater understanding of factors that control the microbial community and microbial processes is needed to achieve the potential of microbial management in agricultural systems. Both biotic and abiotic factors and their interactions may control microbial community composition (6–11), but our ability to predict the key factors and their magnitude of influence in the microbial community structure and functions is limited. Understanding the factors that control plant-associated microbiomes could offer a novel opportunity to engineer microbiomes to support microbiome-based agriculture.

Plant species and genotypes influence root microbiomes, regardless of soil type or geographic location (12–20). However, the effect of plant species or genotype on the microbiome is generally smaller than the effect of environmental and edaphic factors (7, 14, 21, 22). Despite the relatively small magnitude of effect, plant genotypic effects on microbiome composition are particularly important because they indicate the potential for harvesting benefits from microbes indirectly through the choice of crop genotypes. Plant genotypes with desired phenotypes can be used as an engineering tool to select candidate taxa (23, 24). For example, microbial assemblages that are directly associated with high-yielding genotypes may represent candidate taxa for designing microbial consortia with a potential to serve as biofertilizers or biocontrol (25), while exploring the host genes associated with microbial selection may provide insight to support microbiome-focused crop breeding.

Compared to aerial plant surfaces, roots are particularly important for microbe-microbe and host-microbe interactions (14). The endosphere and rhizosphere are active in the exchange of nutrients and microbes between soil and plant (13, 17, 19, 22, 26–28). Microbial diversity and community composition often differ between the rhizosphere and endosphere (13, 17, 19, 22, 29). Models of microbial acquisition in plant root posit a gradual enrichment of microbial communities in the rhizosphere and rhizoplane, followed by pronounced exclusions of microbial communities in the endosphere (29, 30). It seems that the endosphere is more directly under host control and serves as a stronger filter than the rhizosphere, where control is less direct and perhaps driven by chemical cues or abiotic filters along with trophic and non-trophic microbial interactions (29).

Evaluation of the microbial diversity in designed experiments provides an avenue to generate hypotheses about the mechanisms of treatment effects on host phenotype and performance (31). Under the insurance hypothesis, systems with higher species diversity may be more likely to maintain community functions during perturbation (e.g. 32)). Additionally, greater microbial diversity may increase host performance and system robustness. For instance, a host plant lacking resistance to a pathogen under sterile experimental conditions gained the disease-resistant phenotype through the introduction of a phyllosphere microbiome (33), highlighting the potential importance of microbial diversity for host phenotypes/traits (24, 34). In some cases, microbiome composition varied with the level of pathogen infection (34, 35), and particular microbial taxa may be “driver microbes” or “passenger microbes”, as reflected by the magnitude of their effect on host performance (36). In agricultural systems, an increase in overall microbial diversity has been reported in disease suppressive soils, even though few key microbes have been deemed crucial in regulating suppression (37, 38). Greater diversity along with key functional species might drive some ecological functions and failure to select for microbial associates that are antagonistic to pathogens could result in lower host fitness (39, 40). Together these findings indicate that microbiomes can modulate host traits and disease phenotypes and suggest the potential to design agricultural practices to manipulate microbiomes in microbiome-focused crop-improvement programs (23, 24).

Our project addresses central questions about how grafting and the choice of tomato genotype may affect the tomato rhizobiome. Although tomato grafting is a fairly new practice among farmers in the USA, it is an ancient propagation practice in agriculture, commonly used in vegetable production. In the case of tomato, interspecific rootstocks, where rootstock and scion belong to different species of *Solanum,* generally have a rootstock resistant to soilborne diseases (e.g., Fusarium wilt, Verticillium wilt, bacterial wilt, and root-knot nematodes) grafted with a scion that produces higher quality fruit (41–43). Soilborne pathogens are difficult to manage and can result in up to 90% yield loss (44). In addition to the effectiveness of grafted plants in managing soilborne diseases, grafted plants are often more vigorous and more efficient in nutrient uptake and resource utilization, as well as resistant to abiotic stresses (45–48). Thus, plants grafted with effective and vigorous rootstocks often provide higher fruit yield and plant biomass (49, 50). These phenotypic traits of grafted plants appear to be influenced in part by modification of the scion through migration of molecules such as proteins, mRNA and small RNA from rootstocks to scion (51–53), or by epigenetic modifications (54, 55). Restriction of pathogen migration in resistant rootstocks due to pit membrane architecture (56) and the mobility of nucleic acids and proteins (51, 52) are among the physiological and molecular responses of grafted plants during infection. In our study we address how hybrid rootstocks affect the tomato rhizobiome, and directly assess the role of rootstock genotypes and grafting (or artificial selection/breeding) on the rhizobiome. Given the great potential of microbes in plant health and production, and the economic value of grafted tomato, understanding how grafting and rootstock treatments modulate root-associated microbial diversity and community composition will lay the groundwork for future microbiome-based systems with rootstocks supporting higher plant biomass and fruit yield. Further, identification of microbes associated with desirable host traits is a first step to select candidates for synthetic microbiomes.

In this study, we evaluated the effects of rootstocks on tomato rhizobiomes in Midwestern (Kansas, USA) growing conditions. We characterized and compared the composition and diversity of bacterial communities associated with tomato rootstocks by sampling the rhizobiome - microbes within roots (in the endosphere) and surrounding the root (in the rhizosphere). Based on the previous studies of microbial communities in related systems that we review above, our expectation was that : (i) effective rootstocks, defined in terms of fruit yield and plant biomass, will be associated with higher microbial diversity, (ii) differences in microbiome composition will be greater in the endosphere than in the rhizosphere, (iii) the number of taxa responsive to rootstock genotypes (OTUs whose proportion is different than in the nongraft and selfgraft controls) will correlate with rootstock performance, and (iv) responsive taxa will be more frequent in the endosphere than in the rhizosphere. In total, we address the effect of: (i) an agricultural practice (grafting), (ii) rootstock genotypes, (iii) farm sites, and (iv) root compartment (rhizosphere or endosphere) on microbial community composition and diversity.

## MATERIALS AND METHODS

### Rootstocks and experimental design

Plants sampled for this analysis were part of a larger study of grafted tomato plant yield. The main objectives of the larger study were to identify rootstocks that are more productive in Midwestern (Kansas, USA) growing conditions, and to evaluate the effects of rootstocks on the rhizobiome (57). Plants sampled for rhizobiome analyses included three rootstock genotypes (BHN589, RST-04-106, and Maxifort) representing four different treatments: 1) nongrafted BHN589 plants; 2) self-grafted BHN589 plants (plants grafted to their own rootstock); 3) BHN589 grafted on RST-04-106; and 4) BHN589 grafted on Maxifort. The choice of BHN589 as scion was primarily based on the popularity of BHN589 for its high fruit yield and quality and long shelf life. We selected Maxifort because it is a common and popular rootstock, and RST-04-106 as a new rootstock variety based on breeders’ recommendations. In the initial field trials that evaluated rootstock performance, plants grafted with Maxifort had higher yield and biomass, whereas the performance of plants grafted with RST-04-106 was similar to the nongraft and selfgraft controls (Meyer 2016). More information about these rootstocks and their potential disease resistance profiles is available in a USDA resource database (http://www.vegetablegrafting.org/resources/rootstock-tables/tomato-rootstock-table/), and also listed in Table S3.

We established field trials at the Olathe Horticulture Research and Extension Center (OHREC) and in collaboration with local farmers at Common Harvest Farm and Gieringer’s Orchard (Table 1). At each of three study sites the four graft treatments were assigned to four plots per block in a randomized complete block design. Each plot consisted of 5-8 plants, and one of the middle plants from each plot was sampled for the rhizobiome characterization. The number of blocks varied from one study site to another depending on the area available for growing tomato. There were six blocks at the OHREC and four at each of Gieringer’s Farm and Common Harvest Farm, such that for each year, all treatments were replicated 14 times. The experiments were conducted in 2014 and 2015 with similar experimental designs and management. However, at each study site, the blocks were randomly assigned separately for each of the two years.

**TABLE 1.** Sites included in the study, with geographic coordinates and soil type.

| Study sites | Location | Latitude | Longitude | Soil type |
| --- | --- | --- | --- | --- |
| Olathe Horticulture Research and Extension Center (OHREC) | Johnson County, KS | 38.88N | 94.99W | Chase silt loam |
| Common Harvest | Douglas County, KS | 38.96N | 95.20W | Eudora-Kimo complex |
| Gieringer's Orchard | Johnson County, KS | 38.79N | 95.04W | Sibleyville loam |

### Tomato grafting and high tunnel production

Tomato plants were grafted in-house using a tube-grafting technique. Details about the grafting and post-grafting management are available in Meyer (2016). Briefly, young seedlings (9-12 days old) of the scion and the rootstocks were grafted and kept in a healing chamber (dark with high relative humidity: 85-95%) for an additional seven days to facilitate the healing process. Darkness minimizes photosynthesis and high humidity prevents the scion from wilting by maintaining sufficient turgor pressure. Healed plants were transferred to full sunlight in a greenhouse for a week and then transplanted to high tunnels.

All our experiments were conducted in high tunnels, a popular system for tomato production. A high tunnel is an unheated greenhouse, covered with plastic or acrylic glass (plexiglass) with partial ventilation on the sides, and is a relatively new production system among small-scale farmers in the Midwest. High-tunnel production systems protect crops against biotic and abiotic stresses, extend the growing season, and improve fruit quality and yield (58). Details about the high tunnel design and management used in our experiments are provided by Meyer (2016).

### Endosphere and rhizosphere sample preparation

To evaluate microbial communities at the most productive phase of tomato plant development, we sampled during the peak harvest. From each plot, one of the middle plants was carefully dug such that the majority of roots were intact. The intact root mass was shaken ten times to dislodge bulk soil and placed on top of a marked-sampling grid to randomly select four roots, each about 10-12 cm in length (Fig. 1). In most plants, we sampled mainly secondary roots, with few primary roots. In other words, we sampled only roots ~ 1-2 mm in diameter, as these higher order lateral roots are the active organs of nutrient absorption, exudation, and bacterial colonization (59–61). Each root piece was transferred to an individual sterile 15 ml falcon tube (i.e., one piece per tube, and four tubes per plant) containing 10 ml 0.1% Triton-X solution, stored on ice in a cooler, and transported back to the lab within 6 hours, then stored overnight at 4°C. On the following day, roots were sonicated for 10 minutes at high intensity in an ultrasonic bath cleaner (Fisher Scientific, Waltham, MA, USA). High intensity sonication effectively separates rhizoplane communities from endosphere communities (62); in our case the separated rhizoplane communities were included in the rhizosphere samples. The sonicated roots were pressed between sheets of Kimtech Kimwipes (Kimberly-Clark, Roswell, GA, USA), dried, and stored in a 1.5 ml RINO screw-cap tube (Next Advance, Averill Park, NY, USA) at -20° C for DNA extraction. The buffer solution containing the dislodged rhizosphere material was collected into a sterile 20 ml BD Luer Lock disposable syringe (Becton, Dickinson and Company, NJ, USA), and passed through a 0.2 μm Whatman Nuclepore Polycarbonate Track-Etched Membrane filter with 25 mm diameter (Fisher Scientific, Waltham, MA, USA) to collect bacteria and suspended particles. The membrane filters containing rhizosphere contents were stored in a 1.5 ml RINO screw-cap tube at -20° C until DNA extraction. The four root pieces per plant were processed individually until the secondary PCR amplification, and pooled in a single unit prior to sequencing. The Luer Lock filter-holders were cleaned with 10% bleach, rinsed with deionized water, and autoclaved after each run to prevent cross-contamination between samples.

**FIG 1.**
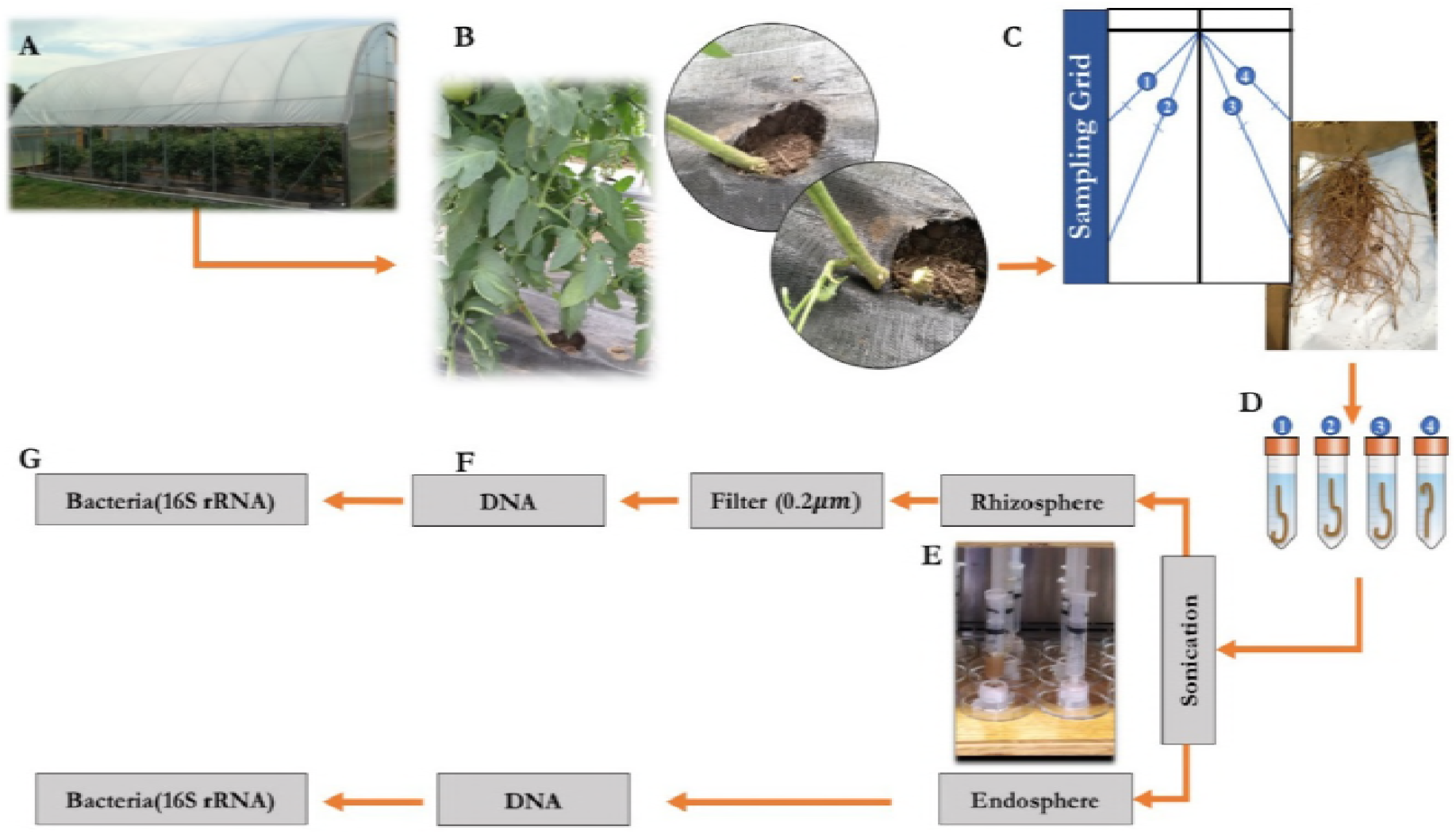
Flowchart of methods for sampling the rhizobiome. Tomato plants grown in high-tunnels (A) were trimmed at the graft-line (B) and four pieces of roots (1-2 mm in diameter) were sampled systematically using a marked-sampling grid (C). Each root piece was transferred to a sterile 15 ml falcon tube containing 10 ml 0.1% Triton-X solution (D), sonicated, and filtered through 0.2 μm filter using a luer filter-holder and disposable syringe (E). DNA was extracted from the endosphere and the rhizosphere samples (F) using an extraction kit, and amplicon libraries representing the bacterial community were prepared using primer sets for the V4 region of 16S rRNA (G).

### DNA extraction and amplicon generation

Total genomic DNA was extracted from both the rhizosphere and root tissue samples using a MoBio Ultra clean soil DNA extraction kit (MoBio, Carlsbad, CA, USA) as per manufacturer’s protocol, with slight modification during the homogenization step. Due to the toughness of the tomato roots, they were cut into smaller pieces using a sterile razor blade, and then homogenized using stainless steel beads in a Bullet Blender (Next Advance, Averill Park, NY, USA) at 4° C. First, we ran a dry homogenization for 10 minutes, and then a wet homogenization for another 10 minutes in IRS and bead solutions from the extraction kit. After extraction, DNA was quantified using a ND1000 spectrophotometer (NanoDrop Technologies, Wilmington, DE, USA) and normalized to 2 ng/ul. Amplicons for the variable region V4 within the bacterial rRNA gene were generated using primers: 515F - GTGCCAGCMGCCGCGGTAA and 806R-GGACTACHVGGGTWTCTAAT (e.g. 63) . Prior to sequencing, amplicons from different plant samples were multiplexed by incorporating unique Molecular Identifier Tags (MIDs) at the 5’ end of the reverse primer.

PCR amplicons were generated in 50 μL reactions under the following conditions: 1 μM forward and reverse primers, 10 ng template DNA, 25 μL 5X Phusion HF Buffer (Finnzymes, Vantaa, Finland) containing 200 μM of each deoxynucleotide and 1.5 mM MgCl_2_ in a mastermix, 15 μL molecular biology grade water, and 1 unit (0.5 μL) Phusion High-Fidelity DNA Polymerase (Finnzymes, Vantaa, Finland). PCR cycle conditions consisted of a 94°C initial denaturing step for 3 minutes, followed by 30 cycles at 94°C for 45 seconds, 50°C annealing for 1 minute, and a 72°C extension for 1.5 minutes, followed by a final extension at 72°C for 10 minutes. To incorporate MID into the PCR amplicons, secondary PCR was run using similar reagents and PCR cycling conditions as in the primary PCR, with the amplification cycles reduced to 10. All DNA samples were amplified in triplicate to minimize PCR stochasticity, pooled, and cleaned using Diffinity RapidTip (Diffinity Genomics, West Chester, PA). Similarly, the secondary PCR was run in triplicate, pooled by experimental unit, and cleaned with Agencourt AmPure cleanup kit using a SPRIplate 96-ring magnet (Beckman Coulter, Beverly, Massachusetts, USA) as per the manufacturer’s protocol. Prior to cleaning the secondary amplicons with an Agencourt AmPure, the amplicons of four pieces of root samples from a particular plant (i.e., an experimental unit) were pooled to a single unit. Then, 500 ng of cleaned barcoded PCR amplicons were combined per experimental unit, and the final pool was cleaned again using an Agencourt AmPure cleanup kit as above. Illumina MiSeq adaptors were ligated to the library and paired-end sequenced on a MiSeq Personal Sequencing System (Illumina, San Diego, CA, USA) using a MiSeq Reagent Kit V2 with 500 cycles. The endosphere and the rhizosphere amplicon libraries were sequenced separately in two runs. Adaptor ligation and sequencing were performed at the Integrated Genomics Facility at Kansas State University.

### Bioinformatics and OTU designation

The sequence data were curated using MOTHUR (Version 1.33.3 (64); following steps outlined in the MiSeq Standard Operating Protocol (SOP; www.mothur.org/wiki/MiSeq_SOP). Briefly, the forward and the reverse sequence reads were contiged using a default criteria as specified in the SOP. Any sequence shorter than 250 base pairs or containing ambiguous bases, more than eight homopolymers, more than one mismatch in primer or MIDs or missing MIDs was removed. Barcoded sequences were assigned to experimental units, and fasta and groups files for the endosphere and the rhizosphere libraries were merged and processed together for the remaining steps in MOTHUR. The cleaned sequences were aligned against curated 16S rRNA gene SILVA alignment, chimeric sequences identified using UCHIME (65) were removed, and the remaining sequences were assigned to taxonomic groups using the Naive Bayesian classifier (66) at 60% bootstrap confidence score against the 16S rRNA gene training set (v9) of the Ribosomal Database Project (67). Sequences without known affinities or assigned to mitochondria or chloroplasts were removed. Pairwise distances (less than 0.10) between aligned DNA sequences were used to cluster the sequences into OTUs at 97% similarity using the nearest-neighbor joining algorithm. However, because of the large distance matrix, sequences were split into bins by taxonomy prior to clustering into OTUs. Finally, the clustered OTUs were assigned to consensus taxonomy and used in community analyses. Due to variation in the sequence yield per sample over two years, we analyzed the libraries for each year separately. To minimize the bias resulting from unequal sequence counts per sample, samples within each year were ratified at the sequence frequency of the sample yielding the lowest count (2568 per sample in 2014, and 8885 per sample in 2015).

### Statistical analyses

To evaluate diversity, we calculated Shannon entropy in R (68) using the vegan package (69) implemented as a part of the phyloseq package (70). The observed Shannon entropy was compared among rootstock treatments using a mixed ANOVA model in the lme4 package (71) in R. Rootstock treatments were compared using study sites as a random factor. Differences in the bacterial communities across rootstocks, compartments and study sites were visualized in non-metric multidimensional scaling (NMDS) plots based on the Bray-Curtis dissimilarity matrix. The observed variation in the bacterial community was partitioned using a permutational multivariate analysis of variance (PERMANOVA, at 1000 permutations) using the adonis function in the vegan package. To identify the OTUs that were depleted or enriched as a function of rootstock treatments, differentially abundant OTUs (DAOTUs) were evaluated by fitting a generalized linear model (GLM) with a negative binomial distribution. Likelihood ratio tests and contrast analyses were performed on the fitted GLM to identify DAOTUs. We used OTU counts from selfgrafts and nongrafts as controls, and compared them with other rootstocks in a contrast analysis. All the tests were adjusted to control for the false discovery rate (FDR, p value < 0.05) using the Benjamini-Hochberg method. Similarly, a differential abundance test was performed comparing the controls (selfgraft vs. nongraft) to identify the OTUs responsive to grafting. General community profiles were constructed using OTUs labeled at the phylum level, split by rootstocks and compartment (endosphere or rhizosphere) and visualized in a bar graph.

### Accession number(s)

All sequence data generated in this study will be deposited in the NCBI Sequence Read Archive depository.

## Results

### General bacterial community data description

The final curated dataset consisted of 1,282,843 sequences from the endosphere- and rhizosphere-associated bacterial communities in tomato. The bacterial communities were dominated by Proteobacteria (37.9%), which was the most abundant phylum across all the rootstock treatments and in both the rhizosphere and endosphere (Fig. 2, Fig. 1S, Table S2). Approximately 14.5% of the sequences in the dataset were unclassified at the phylum level. Among the rootstocks, the selfgraft had the highest percentage of Proteobacteria (39.0%). Actinobacteria (16.9%) and Firmicutes (8.9%) were the other dominant phyla observed in the overall community. Firmicutes and Planctomycetes were enriched in the hybrid rootstocks compared to the nongraft and selfgraft, whereas the Bacteriodetes were depleted in the hybrid rootstocks. At the class level, the communities were largely dominated by α-proteobacteria (18.5%). The class Sphingobacteria was found at a lower percentage in the hybrid rootstocks than in the selfgrafts and nongrafts. Bacteria of class γ-proteobacteria (6.9%) were less dominant in the Maxifort rootstock, compared to other rootstocks. Rhizobiales (13.8%) was the most dominant bacterial order in the overall community, including all rootstock treatments and both compartments. Bacillales were more frequent in Maxifort (7.2%) than in RST-04-106 (6.8%), selfgraft (6.4%), and nongraft (6.2%). At the family level, Planctomycetaceae were more frequent in Maxifort and RST-04-106 (7.4%) than in nongraft (6.9%) and selfgraft (7.0%) treatments. Taxa in the order Myxococcales were more frequent in Maxifort (2.6%) than in the other rootstocks. Analysis at the genus level revealed *Pasturia* spp. as the most dominant taxa in the overall community, as well as in Maxifort. Comparison of community profiles of bacteria between the endosphere and rhizosphere showed that Proteobacteria, Actinobacteria, and Bacteriodetes were more abundant in the endosphere than in the rhizosphere, whereas Planctomycetes, Firmicutes, and TM7 were more abundant in the rhizosphere.

**FIG 2.**
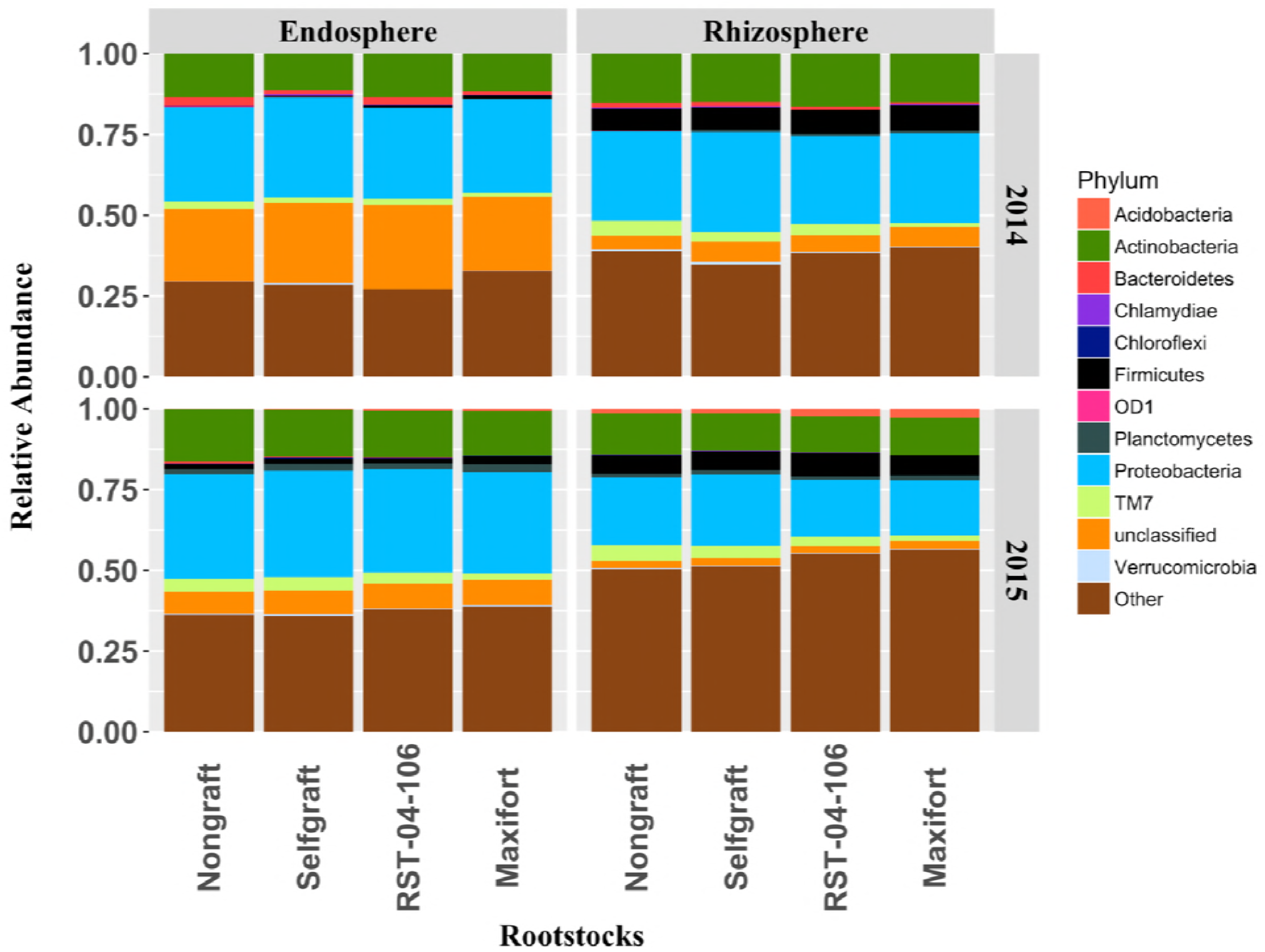
Profile of bacterial communities from tomato at the phylum level. The relative abundance of bacterial phyla recovered from the endosphere and rhizosphere from two hybrid rootstocks (RST-04-106 and Maxifort) and nongraft and selfgraft controls (BHN589) for the years 2014 and 2015. The colored area of each bar represents the relative abundance of the corresponding phylum. The vertical facet in the graph represents root compartments, and the horizontal facet divides the plot by year. OTUs with relative sequence abundance below 1% are summed as “other”.

### Effects of grafting and rootstock on α-diversity

There was strong evidence for a rootstock treatment effect on the diversity of bacteria in the endosphere and rhizosphere compartments of tomato plants (p < 0.01). Among the four treatments, plants grafted with the Maxifort rootstock had the highest diversity compared to selfgrafted and nongrafted plants in both the endosphere and the rhizosphere in 2014 (p < 0.05; Fig. 3). In 2015 we observed similar results with the highest diversity in the Maxifort-grafted plants for both the compartments; however, there was less evidence (p = 0.1) for a difference between the selfgraft and the Maxifort-grafted treatments in the endosphere (Fig. 3). The effect of RST-04-106 on bacterial diversity was similar to that of the nongraft and selfgraft treatments (p > 0.05). Additionally, there was not evidence for a difference between the selfgrafted and the nongrafted plants in the endosphere (p = 0.9) or in the rhizosphere (p = 0.2). When the effect of rootstock treatments (a fixed effect) on bacterial diversity was evaluated for each compartment, study site, and year separately in a factorial design, the effects of rootstocks varied (Table S1), though there was never strong evidence for an interaction between rootstock treatments and study sites (p > 0.05).

**FIG 3.**
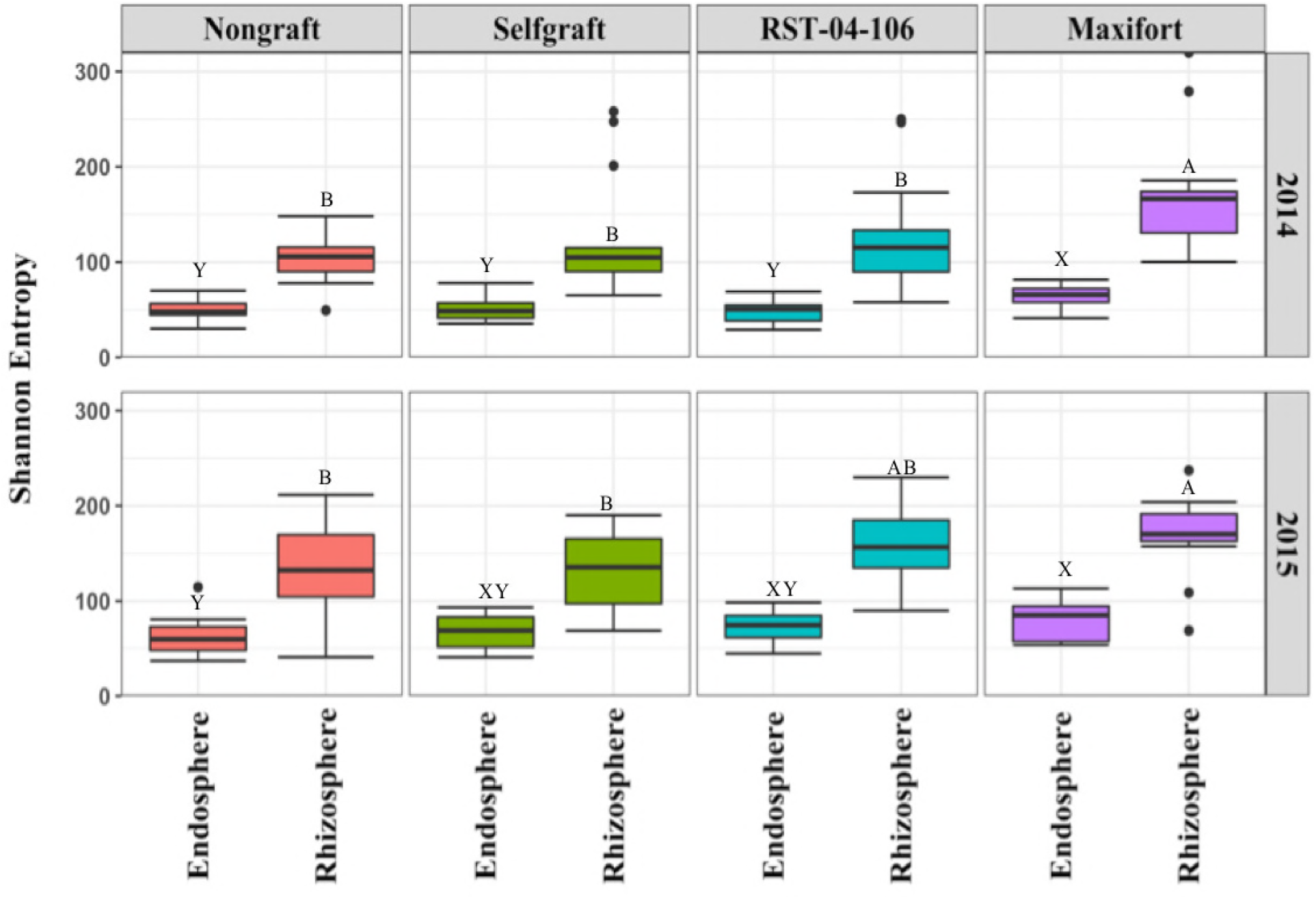
Diversity of bacterial communities associated with tomato. Bacterial diversity was evaluated for the endosphere and rhizosphere from two hybrid rootstocks (RST-04-106 and Maxifort) and nongraft and selfgraft controls (BHN589) in a mixed model ANOVA. Rootstock treatments were compared using study site as a random factor. The plot is vertically faceted by rootstock treatments, and the horizontal facets represent sample years. Shannon entropy (a measure of α-diversity) was higher for Maxifort compared to the nongrafted controls (p < 0.05). There was strong evidence that Shannon entropy differed for treatment combinations with different letters (p < 0.05).

### Effects of grafting and rootstock on bacterial composition

PERMANOVA analysis based on the Bray-Curtis dissimilarity distance matrix identified rootstock treatment, study site, and endosphere-rhizosphere compartments as factors explaining the variation in the bacterial community. The percentage of variation explained by the rootstock treatment was small (3%) but consistent across the two years (PERMANOVA in year 2014: df = 3, *F_model_* = 2.0, R^2^ = 0.03, *P* = 0.02; PERMANOVA in year 2015: df = 3, *F_model_* = 1.9, R^2^ = 0.03, *P* = 0.01). Visualization of the distance matrix for samples using NMDS ordination plots (Fig. 4) suggested overlapping centroids across the rootstock genotypes, whereas compartment and study site were the primary factors partitioning communities in both years. Endosphere-rhizosphere compartments (PERMANOVA in year 2014: df = 1, *F_model_* = 58, R^2^ = 0.28, *P* = 0.001; PERMANOVA in year 2015: df = 1, *F_model_* = 53, R^2^ = 0.25, *P* = 0.001) and study site (PERMANOVA in year 2014: df = 2, *F_model_* = 18, R^2^ = 0.17, *P* = 0.001; PERMANOVA in year 2015: df = 2, *F_model_* = 21, R^2^ = 0.20, *P* = 0.001) explained about 45% of the variation in the bacterial community.

**FIG 4.**
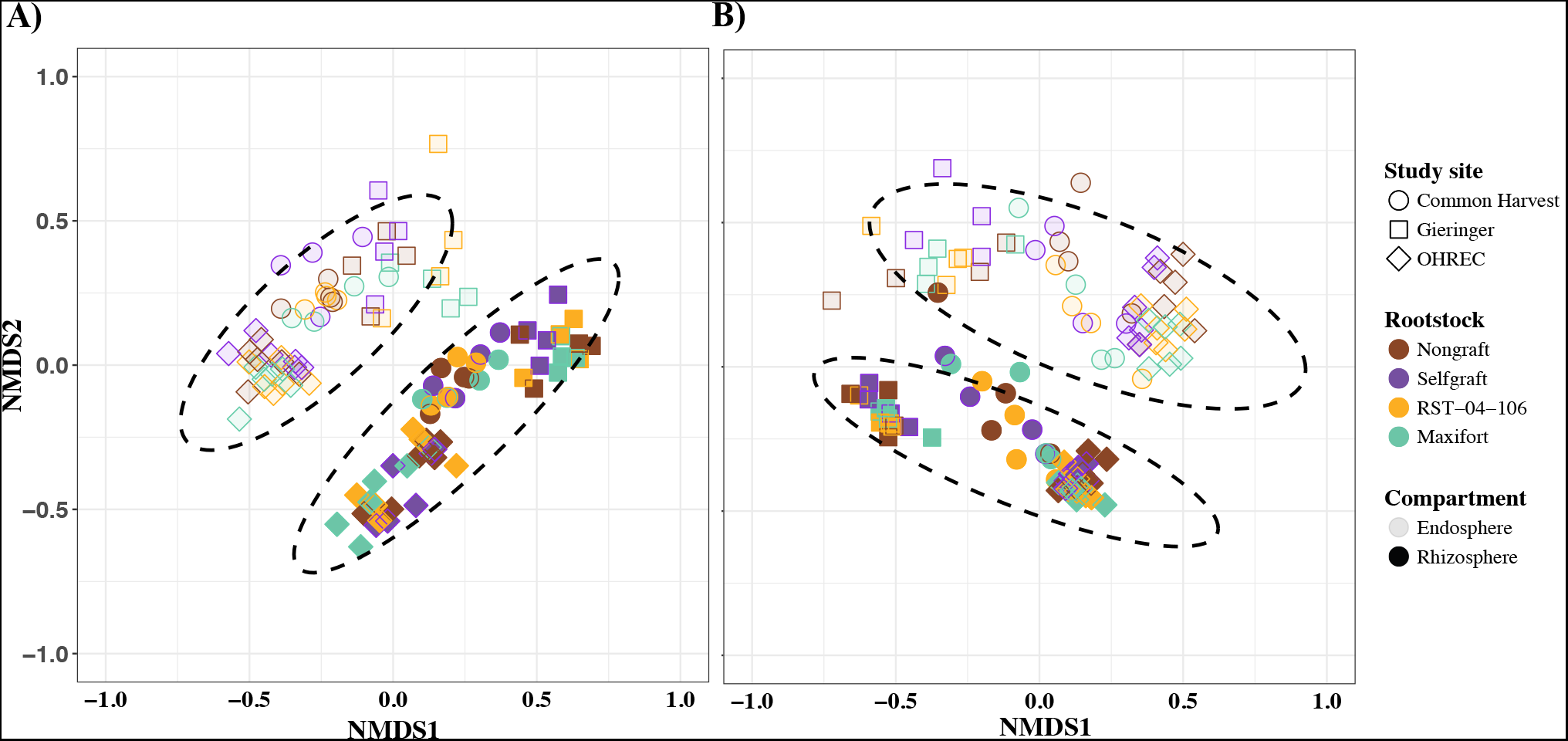
Non-metric multidimensional scaling (NMDS) ordination of samples from tomato rootstock treatments. NMDS ordination of samples based on the Bray-Curtis dissimilarity distance matrix of OTUs from bacterial communities inhabiting the endosphere and rhizosphere compartments in years 2014 (A) and 2015 (B). Color indicates rootstock treatment (two hybrid rootstocks (RST-04-106 and Maxifort) and nongraft and selfgraft controls (BHN589)), shape represents study site, and solid and lighter fill color represents the rhizosphere and endosphere compartment respectively. Ellipses indicate a 95% confidence interval around the centroid of endosphere and rhizosphere samples.

### Comparison of DAOTUs

Bacterial diversity and community composition in the tomato endosphere and rhizosphere differed among rootstock treatments. To identify taxa that responded to rootstocks, we used a differential abundance test. Although we consistently observed higher α-diversity in the rhizosphere than in the endosphere, the total number of DAOTUs, either enriched or depleted, was greater in the endosphere (n=56) than in the rhizosphere (n=15; Fig. 5). The analysis of contrasts designed to compare OTU proportion to controls (nongrafts and selfgrafts) found a higher number of responsive taxa in Maxifort in both the endosphere (n=41) and rhizosphere (n=13). Enriched OTUs in the Maxifort rhizosphere included OTUs assigned to the following taxa: Gp5 and Gp6 within phylum Actinobacteria, *Ohtaekwangnia* sp. in Bacteriodetes, *Leuconostoc* sp. in Firmicutes, and three unclassified OTUs (Fig. 5). In contrast, depleted OTUs in the rhizosphere included OTUs assigned to following taxa: two in the TM7 group, two in Proteobacteria, and two that remained unclassified. OTUs such as *Methlophaga* sp. (Proteobacteria), *Blastopirellula* sp. (Planctomycetes), *Halocella* sp. (Firmicutes), *Opitutus* sp. (Verrucomicrobia), Gp16 (Acidobacteria), Gp6 (Acidobacteria), and *Steroidobacter* sp. (Proteobacteria) were enriched in the Maxifort endosphere. In contrast, OTUs with taxonomic affinities to *Sphingobacterium* sp. (Bacteroidetes), *Halomonas* sp. (Proteobacteria), *Chryseobacterium* sp. (Bacteroidetes), *Shigella* sp. (Proteobacteria) and *Flavobacterium* sp. (Bacteroidetes) were depleted in the Maxifort endosphere. A total of eight OTUs changed significantly compared to the controls in the endosphere community of RST-04-106, where six OTUs were enriched and two were depleted. OTUs enriched in the RST-04-106 endosphere included *Spartobacteria* sp. (Verrucomicrobia), and five OTUs unclassified at the genus level, whereas the proportion of *Flavobacterium* sp. (Bacteroidetes) was significantly reduced. The RST-04-106 rhizosphere community was resilient to the grafting treatment; only two OTUs were enriched compared to the controls. Enriched OTUs in the rhizosphere belonged to *Methylocaldum* sp. (Proteobacteria), and an unclassified OTU in phylum Actinobacteria. The bacterial communities in the selfgraft and nongraft controls were surprisingly similar in the rhizosphere – none of the OTUs changed significantly. Although no effect was observed in the rhizosphere, an effect of grafting was observed in the endosphere community, where the proportion of six OTUs changed compared to nongraft control, out of which one (unclassified) was enriched and five were depleted (Fig. 5).

**FIG 5.**
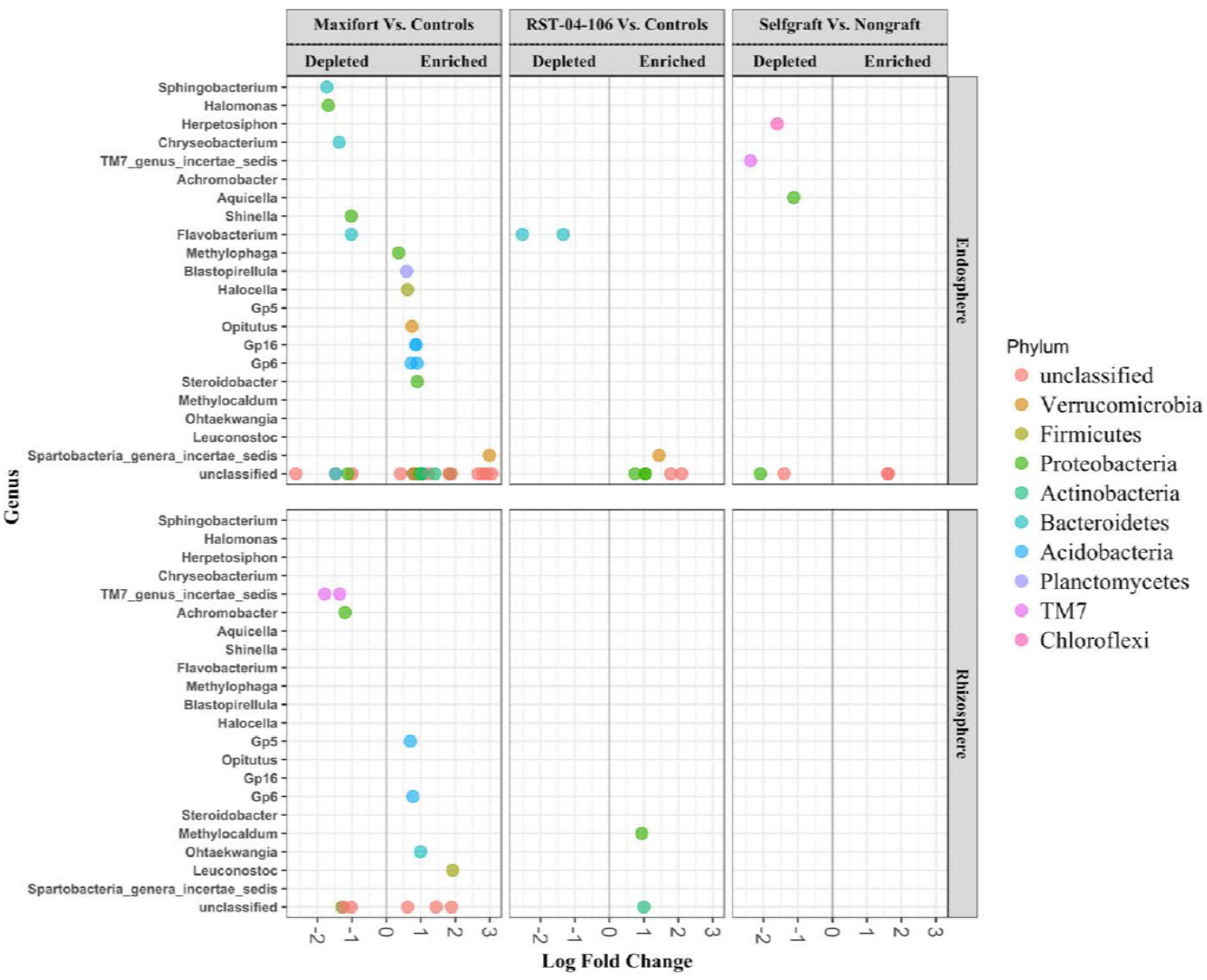
Comparisons of differentially abundant OTUs. Differentially abundant OTUs (DAOTUs) across tomato rootstocks evaluated for the endosphere and the rhizosphere, with OTU counts from selfgrafts and nongrafts (BHN589) as controls. All the tests were adjusted to control for the false discovery rate (FDR, p value < 0.05) using the Benjamini-Hochberg method. Each point represents an OTU labeled at the genus level and colored based on the associated phylum. The position along the x-axis represents the frequency fold-change contrasted with the two controls (selfgraft and nongraft (BHN589)) for the two hybrid rootstocks (Maxifort and RST-04-106), and a selfgraft vs. nongraft (BHN589) contrast.

## Discussion

Rootstocks had an effect on microbial diversity and community composition. We observed higher bacterial diversity in the endosphere and rhizosphere of the high-yielding Maxifort rootstock, compared to nongraft and selfgraft controls of BHN589. Several studies have reported host genotypic effects on microbiome composition; the effects of host genotype on microbiome composition remain poorly understood, but the host effect has generally been smaller than the effect of environmental and edaphic factors (7, 72). Consistent with these published results, tomato rootstocks explained only roughly 3% of the compositional variation in the bacterial community, whereas there was strong evidence that the compartment (endosphere vs. rhizosphere) and the study site explained a major portion of the variation (25-28% and 17-20%, respectively). The magnitude of the effect of host genotypes has been similar in other studies. For example, a study evaluating six different rice genotypes found that only 1.5-2.5% of community composition variation was attributable to rice genotype (22). However, it is important to keep in mind that variation in taxonomic composition may not be strongly reflective of variation in function; plant genotypes may have substantial effects specifically on microbial taxa that are particularly important to plant health.

Differential abundance tests of OTUs across treatments found a greater number of responsive taxa in Maxifort than in the other rootstocks. Interestingly, the endosphere had a greater number of responsive OTUs than the rhizosphere, while α-diversity was higher in the rhizosphere, corroborating the results from grafting studies with grapevine rootstocks (72). We expected that hybrid rootstocks would have higher diversity as well as more DAOTUs than the control treatments, and this was the case for Maxifort compared to the nongraft control (BHN589), but not for the other hybrid rootstock, RST-04-106. Note that the RST-04-106 rootstock also had lower performance in terms of yield and biomass in our field trial (57), consistent with the idea that bacterial diversity and host performance may be associated with graft performance.

Earlier agricultural microbiome studies have primarily focused on how a particular management strategy – such as organic versus conventional farming (73, 74) or tillage practices (75, 76) – influence soil microbial communities, and rarely incorporate host genotypic effects and their interaction with agricultural practices. Using tomato as a grafting model, our study provides a new perspective on the effects of host genotype on microbial communities. Our results for microbial diversity and composition suggest that grafting with specific rootstocks influences microbiome assembly as well as yield and biomass. Our studies included only two hybrid rootstocks and one scion (BHN589), so research including more rootstocks and scions along with hybrid rootstock specific selfgraft and nongraft controls would help to generalize these results. Given the economic importance of grafted tomatoes, and the need to develop sustainable production systems, our findings indicate new opportunities for improving microbiome-based practices in agriculture.

The rootstocks included in our study represent different genetic backgrounds and are specifically bred to provide resistance to multiple soilborne diseases (Table S3). Variation in a plant genetic background influences the acquisition of root-associated microbial communities (77), as shown in rice (22), maize (18), and *Arabidopsis* (13, 17, 78). Physiologically, hosts vary in their root exudates and rhizodeposits (79–81), thereby creating host specific cues to select microbial associates from the surrounding soil (29, 82, 83). Root exudates and rhizodeposits not only contain carbon and other nutrients that support the belowground food web, but also contain chemicals such as cytokinins (84), phytotoxins, antibiotics, and hormones (85) that are key to supporting some microbial assemblages while deterring others (86, 87). These findings indicate an active and host-specific microbiome filter, although the extent of such selection may differ across plant types and studies (17, 19, 21, 22, 24, 27). Usually, when plant types are closely related, their microbial communities are more similar (29), and smaller differences are observed in selection of microbial communities by plant type (88).

The size of the effect of tomato rootstocks on rhizobiome diversity and composition depends on which root compartment is being considered. The tomato rhizosphere supported a more diverse microbial community than the endosphere. These results were consistent with other studies of plant-based selection of microbial communities, and with a proposed model of microbial acquisition (29), where the endosphere microbial community is under more direct host control (89, 90) and is more strongly filtered than the rhizosphere, where control is less direct and may be driven by chemical cues or inter-microbe interactions (59). These results confirm the effect of agricultural practices on the acquisition of endosphere and rhizosphere microbiomes, and are particularly interesting from a practical standpoint. Microbes in the rhizosphere microbiome could represent candidate taxa for biofertilizers or plant supplements, and understanding the mechanisms underlying the host-based selection of endosphere microbiome members could guide microbiome-based breeding programs. While research efforts to support such breeding programs are still limited, plant loci identified for host-genotype dependent structuring of microbial community via GWAS (91) are a promising first step.

Root architecture and anatomy change to mediate plant responses to biotic and abiotic stresses (92–94), and can vary within and between plant species and genotypes (59, 95). If we learn the extent to which host genotypes affect root architecture and physiology, such as exudation, then we may also understand the potential mechanisms by which host genotypes control microbiomes via metabolites and exudates (82, 83). Genotypic variation in root traits and functions selects for different microbial communities, by modulating the quality as well as the quantity of root exudates and by modifying the physical and chemical properties of the surrounding soil environment (96). For instance, a greater root mass with more abundant fine roots was reported for Maxifort, along with greater arbuscular mycorrhizal colonization (97). We did not evaluate root architecture in our experiments, but future studies of microbial diversity for root types (e.g. primary roots, secondary roots, root hairs) and the effects of root exudates will help to clarify the process of microbial acquisition by rootstock genotypes.

Previous studies in our experimental system (57) and other grafted tomato systems (50, 98, 99) have indicated that tomatoes with effective rootstocks gain greater aboveground biomass and have higher overall photosynthetic activity, a result often attributed to the root system. This aboveground gain increases the total leaf area, likely resulting in a greater supply of photosynthates belowground. Translocation of fixed carbon from shoots to roots may allow plant roots to actively recruit and sustain diverse microbiomes (100). It is generally accepted that 30-60% of plant photosynthate is transported belowground, much of which (40-90%) is excreted into the rhizosphere, supporting diverse root-associated microorganisms (101, 102). Thus, the higher microbial diversity observed in the rhizosphere of plants grafted with the Maxifort rootstock may be a result of its vigorous root system (97), resulting in greater shoot biomass as well. Grafting may provide a boost in the continuous feedback between the aboveground and belowground compartments, and consequently greater investment of the photosynthetic capital in recruiting diverse microbial communities.

Using differential abundance tests, we identified OTUs that were sensitive to the rootstock treatments. These OTUs were either enriched or depleted in contrast to the controls, and potentially represent taxa that were under direct selection by rootstocks and their chemical cues. We expected there would be a higher number/percentage of responsive OTUs (or DAOTUs) in the endosphere than in the rhizosphere, and our results support this, corroborating results from other studies (22, 72). More DAOTUs in the endosphere relative to the rhizosphere might be a consequence of direct host control in the endosphere microbiome (103, 104). In addition to the compartment-specific effect on the number of sensitive taxa, it was interesting to observe the higher number of sensitive OTUs in Maxifort compared to the other treatments. The results are consistent with the expectation that enhanced host performance is associated with an increase in responsive taxa. Responsive taxa could be important for host performance, as differentially abundant taxa are often functionally associated with host physiology and immune system (78, 105). However, many of the responsive taxa in this study are unclassified, and may be unculturable. Future experiments are needed to evaluate the influence of DAOTUs on host phenotypes.

Taxa that were enriched in Maxifort included representatives of Firmicutes, Verrucomicrobia, Planctomycetes, Proteobacteria and Acidobacteria. We observed significant depletion of TM7, also known as *Candidatus Saccharibacteria,* in the rhizosphere of Maxifort. TM7, previously reported as an antagonist to the antibiotic-producing Actinobacteria (106), can also be involved in suppressing host immune systems, and causing disease (107–109). The observed proportion of Actinomycetes was two-fold higher in both the Maxifort endosphere and rhizosphere compared to TM7 (p < 0.001 in both cases, Mann-Whitney-Wilcoxon test). It is possible that the productivity of Maxifort is due, in part, to selection for antibiotic-producing bacteria that are detrimental to pathogenic bacteria. Further studies with taxa identified in this study would be necessary to evaluate this hypothesis, especially in the presence of the relevant pathogens. Note that we did not observe any obvious disease symptoms in our experiments. Under disease pressure, greater effects of rootstock treatments on g microbiomes would be likely.

The other OTUs enriched for Maxifort in DAOTU analysis included members of Planctomycetes. Members of this phylum are efficient cross-feeders of exopolysaccharides (EPS), commonly found with nitrogen fixing bacteria (110), and in environments rich in organic matter and nitrate (111). In addition, some members of Planctomycetes are highly tolerant of environmental stressors such as seawater, acidic peat bogs, hot springs, and low temperatures (112–114). Among the other phyla that were enriched were Proteobacteria, which include many important plant-growth promoting (PGP) organisms, but note that some of the OTUs from Proteobacteria were depleted as well, especially *Halomonas* sp. and *Shinella* sp. This analysis also identified some oligotrophic groups in Verrucomicrobia that have been reported in association with pre-agricultural tallgrass prairie soil (115). Past studies have found a decline in Verrucomicrobia with nutrient amendments (116), while they were dominant and functionally active in undisturbed soil with recalcitrant carbon compounds (115, 117). Many of rhizobiome taxa in our study currently lack information about biological function, pointing out the need for further development of culturing methods, along with experiments to test biological functions, to support the design of synthetic communities in microbiome-based crop production.

## Conclusions

Multiple factors define the structure and function of plant-associated microbiomes. Understanding these factors and their control of plant microbiome assembly will support future strategies to augment specific microbes for crop production and disease management. Our studies of a grafted tomato system found evidence for a small contribution of rootstocks in determining the microbial community. The effect attributable to plant compartment (endosphere vs. rhizosphere) was 9-10-fold greater, whereas the effect of study site was 6-7 fold greater than the rootstock effect. We also identified microbes specific to rootstock treatments. Further study of select microbes could help to identify candidate taxa for synthetic communities. In the long term, identifying specific plant alleles/genes that correlate with target microbial taxa can inform plant breeding to meet goals beyond fruit traits and yield, perhaps including means to facilitate low-input sustainable agriculture through plant-mediated selection of desirable microbiome components. These observations of how rootstock treatments, environment, and plant compartment (endosphere vs. rhizosphere) can structure microbial communities help to lay the groundwork for the development of designer communities and microbiome-based breeding to improve crop production.

## Acknowledgements

We thank Alison Cioffi, Bryan S. Cordova, and Jeanelle L. Brisbane for their assistance during the sample processing and DNA extraction; Lani Meyer and Kimberly Oxley at OHREC for coordinating the field visit and sampling; and Tom Buller and Jill Elmers of Common Harvest Farms, and Alicia Ellingsworth and Katherine Kelly of Gibbs Road Farm for their collaboration. We are extremely appreciative of support from The Ceres Trust, a USDA NCR SARE Research and Education Grant (LNC13-355), the Kansas Agricultural Experiment Station, the National Institute for Mathematical and Biological Synthesis (NIMBioS), and the University of Florida. Jumpponen was supported in part by USDA-NIFA capacity project KS-495.

K.A.G., A.J., M.M.K., C.L.R., and R.P. designed the study. R.P. and L.G.M. collected and processed the samples. R.P. analyzed and interpreted the data. R.P., K.A.G., and A.J. wrote the manuscript. R.P., K.A.G., A.J., M.M.K., C.L.R., and L.G.M. revised the manuscript. All authors read and approved the final manuscript.

